# CRISPR Mediated Transactivation in the Human Disease Vector *Aedes aegypti*

**DOI:** 10.1101/2022.08.31.505972

**Authors:** Michelle Bui, Elena Dalla Benetta, Yuemei Dong, Yunchong Zhao, Ting Yang, Ming Li, Igor A Antoshechkin, Anna Buchman, Vanessa Bottino-Rojas, Anthony A. James, Michael W. Perry, George Dimopoulos, Omar S Akbari

**Affiliations:** School of Biological Sciences, Department of Cell and Developmental Biology, University of California, San Diego, La Jolla, California 92093, USA; W. Harry Feinstone Department of Molecular Microbiology and Immunology, Bloomberg School of Public Health, Johns Hopkins University, Baltimore, Maryland, USA; Division of Biology and Biological Engineering, California Institute of Technology, Pasadena, California, 91125, USA; Verily Life Sciences, South San Francisco, CA 94080, USA; Department of Microbiology & Molecular Genetics, University of California, Irvine, USA; Department of Molecular Biology & Biochemistry, University of California, Irvine, USA

**Keywords:** CRISPR, mosquitoes, *Aedes aegypti*, transactivation, dengue virus, virus transmission

## Abstract

As a major insect vector of multiple arboviruses, *Aedes aegypti* poses a significant global health and economic burden. A number of genetic engineering tools have been exploited to understand its biology with the goal of reducing its impact. For example, current tools have focused on knocking-down RNA transcripts, inducing loss-of-function mutations or expressing exogenous DNA. However, methods for transactivating endogenous genes have not been developed. To fill this void, here we developed a CRISPR activation (CRISPRa) system in *Ae. aegypti* to transactivate target gene expression. Gene expression is activated through pairing a catalytically-inactive (‘dead’) Cas9 (dCas9) with a highly-active tripartite activator, VP64-p65-Rta (VPR) and synthetic guide RNA (sgRNA) complementary to a user defined target-gene promoter region. As a proof of concept, we demonstrate that engineered *Ae. aegypti* mosquitoes harboring a binary CRISPRa system can be used to effectively overexpress two developmental genes, *even-skipped (eve)* and *hedgehog (hh)*, resulting in observable morphological phenotypes. We also used this system to overexpress the positive transcriptional regulator of the Toll immune pathway known as *AaRel1*, which resulted in a significant suppression of dengue virus serotype 2 (DENV2). This system provides a versatile tool for research pathways not previously possible in *Ae. aegypti*, such as programmed overexpression of endogenous genes, and may lead to the development of innovative vector control tools.

## Introduction

The yellow fever mosquito, *Aedes aegypti*, is a competent vector of arboviruses including chikungunya, dengue, and Zika [1–3]. Their vectorial capacity, desiccation-tolerant eggs, adaptability to a range of climates and anthropophilic behavior have enabled them to become and remain an increasing burden to human welfare [4–6]. As global temperatures increase, the spatial distribution and range of *Ae. aegypti*, as well as the pathogens they transmit, continue to expand [7]. Historically, insecticides have been the major tool for reducing mosquito populations to control the spread of mosquito-borne diseases. However, as mosquito populations continue to thrive, and evolve resistance to insecticides, alternative control measures are of utmost demand. In particular, strategies centered around genetic manipulation have become a major focus for novel genetic-based tool development.

In tandem with improved and expanding assemblies of the *Ae. aegypti* genome and various transcriptomes [8–10], pivotal tools in the development of genetic-based vector control strategies have been reduction of transcript levels using RNAi [11], germline transformation using transposable elements [12], and programmable DNA targeting-using Clustered Regularly Interspaced Short Palindromic Repeat (CRISPR) [13–15]. For example, RNAi and other small RNAs have been instrumental in generating knock-down phenotypes within the mosquito [16–18] as well as a method for reducing viral transcript numbers [19–23]. Moreover, CRISPR/Cas9 has been pivotal for site-directed mutagenesis and site-specific recombination to become more efficient, precise, and accessible [24,25]. For example, CRISPR/Cas9 mutagenesis has been utilized in *Ae. aegypti* for furthering the understanding of various factors of mosquito biology, such as sex determination [26,27], olfaction [28], behavior [29] and development [30–36]. In addition to site-directed mutagenesis, CRISPR/Cas9 has been used to integrate desired DNA sequences into the mosquito genome through Homology Directed Repair (HDR)[37]. In conjunction, these tools have been used to develop numerous vector control strategies in *Ae. aegypti* including those based on conferring dengue and Zika virus resistance [38,39], homing based gene drives [40,41] and recently the precision-guided sterile insect technique (pgSIT) [42]. Although these tools have been instrumental to the expansion of vector control methods, the ability to specifically induce endogenous gene overexpression has been non-existent.

Recently, researchers have engineered a CRISPR-based tool able to function as a programmable transcription factor for transactivating the expression of target genes [43,44]. The system, known as CRISPR activation (CRISPRa), utilizes a nuclease-deactivated, or dead, Cas9 (dCas9) able to bind to a target locus with the aid of a complementary spacer sequence called a small guide RNA (sgRNA) [45,46]. However, unlike Cas9, dCas9 contains two mutations that disable its endonucleolytic activity, thus preventing cleavage of DNA [47]. Interestingly, when fused with the tripartite transcriptional activators VP64-p65-Rta (VPR), dCas9 can efficiently recruit transcriptional machinery to a promoter region by mimicking the natural cooperative recruitment process of transcription initiation [48]. Consequently, dCas9-VPR is able to transactivate the expression of a specific endogenous gene when guided to the promoter region of the gene of choice [49]. Furthermore, dCas9-VPR can be directed to nearly any DNA sequence with a sgRNA, requiring only a short protospacer adjacent motif (PAM) site 5’-NGG-3’ proximal to the target. With the ability to bind and recruit transcription factors, CRISPRa has previously been utilized to effectively upregulate target genes in human cells, *Bombyx mori* cell lines, and in *Drosophila melanogaster* [50]. Interestingly, this tool also has been leveraged to develop synthetic species in *D. melanogaster* [51], a technique that could prove valuable if engineered in other species, for example mosquitoes. However, *in vivo* use of this tool is limited and it has not been demonstrated yet in any vector-borne insect.

Here we have generated the first CRISPRa system in *Ae. aegypti*. To determine the efficacy of our system we targeted the expression of two conserved developmental genes, *even-skipped* (*eve*, AAEL007369) and *hedgehog* (*hh*, AAEL006708) that play instrumental roles in the spatial and temporal control of embryonic developmental patterning. *Eve* is involved in the development of odd- and even-numbered parasegments [52], whereas *hh* signaling plays numerous roles such as the development of segment polarity and various organs [53]. Following transactivation of these genes using our CRISPRa system, we quantified targeting overexpression using both qPCR and RNA sequencing. In addition, we observed phenotypic changes such as lethality and spatial cellular mispatterning visualized by *in situ* hybridization. Similarly, as a proof of principle we applied the CRISPRa system to transactivate the positive transcriptional regulator gene of Toll immune pathway, known as *AaRel1 or Rel1* (AAEL007696). Rel1 plays a central role as a positive transcriptional factor in antiviral and anti-fungal defenses. The activation of the Toll pathway can be monitored through the transcriptional activation of the up-regulation of the *Rel1* transcription factor or down-regulation of the negative regulator *Cactus*. Transgenic overexpression of *AaRel1* or RNA interference (RNAi)-mediated silencing of the negative regulator *Cactus* have demonstrated the potent antiviral and anti-fungal role of the Toll immune pathway [54–57]. Here we have shown the CRISPRa-mediated transactivation of *AaRel1* resulted in a similar level of suppression of viral (DENV2) infection in the *Ae. aegypti* mosquitoes as that has been observed in the *Cactus* gene silenced mosquitoes [56]. Given these results, CRISPRa can be a powerful tool for investigating gene function in *Ae. aegypti* as well as the basis for the development of novel genetics control strategies.

## Results

### Generation of CRISPRa transgenic lines in *Ae. aegypti*

To engineer a CRISPRa system in *Ae. aegypti*, we generated a transgenic mosquito line that expresses dCas9 fused to a transactivator as well as multiple lines expressing sgRNAs targeting select genes (**Fig. 1**). Based on previous CRISPRa systems created in *D. melanogaster [58,59]* we used dCas9-VPR, a catalytically dead Cas9 (dCas9) fused to the transcriptional activator factor VP64-p65-RTA (VPR), as the dCas9 transactivator. Expression of dCas9-VPR was driven by a *polyubiquitin* (PUb) promoter known to encode relatively high expression levels within a variety of cell types and developmental stages [60] (**Fig. 1B**). We generated two transgenic lines expressing sgRNAs targeting either *even-skipped (eve)*, or *hedgehog (hh)*. To increase efficacy, each sgRNA line encodes two sgRNAs targeting two different regions within the promoter region of the gene of interest (**Fig. 1B**). Specifically, the sgRNA target sites were designed from sequences within 0-250 base-pairs (bp) to the 5’-end of the transcription start site (TSS) to enable robust target gene transactivation (**Fig. 1A**). Expression of sgRNAs was ubiquitously driven by either U6a or U6b promoters [61]. Fluorescent markers were encoded to confirm integration of the transgenes into the mosquito genome as well as tracking inheritance of the constructs (**Fig. 1B, C**).

**Figure 1.**
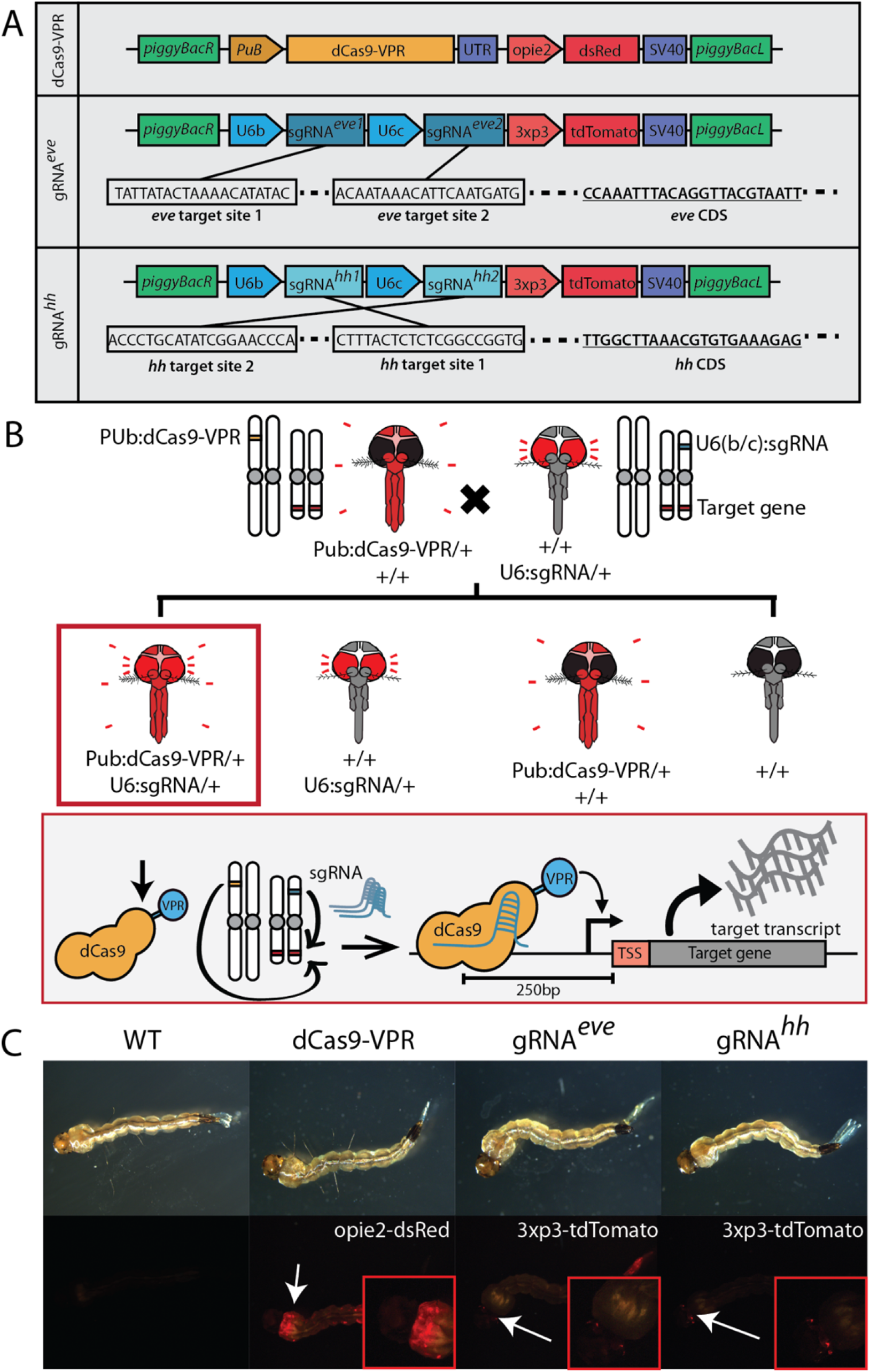
*Ae. aegypti* CRISPRa system transgenic line design and marker expression. **A)** A binary CRISPRa system was designed using two separate transgenic *Ae. aegypti* lines that when crossed, result in transheterozygous individuals expressing both dCas9-VPR and sgRNAs and have upregulated expression of the target gene. **B)** The transgenic lines used in this study include one line expressing dCas9-VPR under a polyubiquitin promoter, a sgRNA line targeting *eve*, and a sgRNA line targeting *hh*. The sgRNA lines were designed to express two distinct sgRNAs targeting the same promoter region of the respective target gene. **C)** dCas9 and sgRNA lines were marked with opie2-dsRed and 3xP3-tdTomato, respectively.

### CRISPRa induces target gene overexpression

To quantify CRISPRa mediated transactivation, we performed genetic crosses between PUb:dCas9-VPR/+ males and U6:sgRNA/+ females. Resulting progeny were collected to quantify expression and transcript abundance of the target genes and observe overexpression phenotypes (**Fig. 1A**). We measured *eve* and *hh* transcript levels using both quantitative real-time PCR (qPCR) and RNA transcriptome sequencing (RNAseq). Total RNA extracted from 24h post oviposition eggs transheterozygous for dCas9-VPR and U6-sgRNA (PUb:dCas9-VPR/U6-sgRNA^*eve*^and PUb:dCas9-VPR/U6-sgRNA^*hh*^), and controls including the two parental lines (PUb:dCas9-VPR; U6-sgRNA^*eve*^and U6-sgRNA^*hh*^), and wild-type controls (Liverpool strain) were all used to measure transcript levels of *eve* and *hh* (15 samples in total). Total RNA samples were divided in two, one sample was used for qPCR analysis and the other half for RNA sequencing. qPCR analysis of *eve* overexpression in transheterozygous individuals (PUB:dCas9-VPR/U6-sgRNA^*eve*^) resulted in a 22.8 (+ 0.75)-fold increase in transcript level compared to wild type and the two controls (F_3,8_ = 659.7; *P* = 6.46e-10; one-way ANOVA and Tukey’s multiple-comparison test) (**Fig. 2A**). Overexpression of *hh* in transheterozygous eggs (PUb:dCas9-VPR/U6-sgRNA^*hh*^) was 8.20 (+ 0.47)-fold higher compared to wild type and the two controls (PUb:dCas9-VPR and U6-sgRNA^*hh*^) (F_3,8_ = 82.67; *P* = 2.32e-06; one-way ANOVA and Tukey’s multiple-comparison test) (**Fig. 2A**).

**Figure 2.**
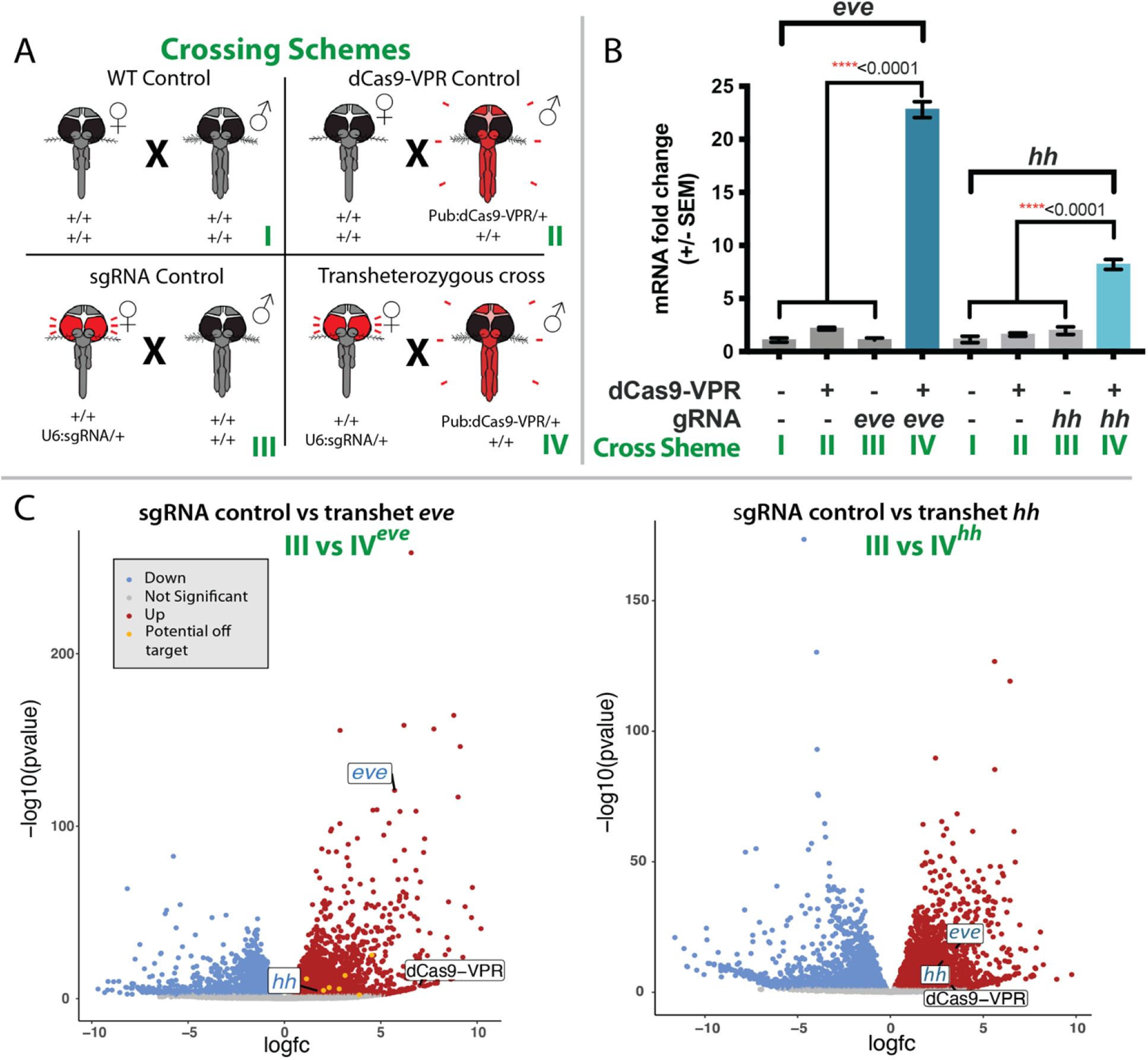
Quantified overexpression of CRISPRa transactivated *eve* and *hh*. **A)** Multiple crosses were performed in order to determine if the CRISPRa system could transactivate target genes. Crosses included a wild-type control (+/+ ♀ X +/+ ♂, I), dCas9-VPR Control (+/+ ♀ X Pub:dCas9-VPR/+ ♂, II), sgRNA control (+/U6:sgRNA ♀ X +/+ ♂, III), and a transheterozygous cross (+/U6:sgRNA ♀ X Pub:dCas9-VPR/+ ♂, IV). **B)** Fold change of expression of targeted genes from CRISPRa transactivation was measured and calculated using qPCR. Differences in mRNA fold change between control crosses and transheterozygous crosses were calculated using a one-way ANOVA and Tukey’s multiple-comparison test, \*\*\*\**p <* 0.0001. **C)** Volcano plots of the RNA sequencing data were to compare expression levels of *eve* and *hh* as well as other genes potentially affected. Down-regulated genes are colored blue while upregulated genes are in red. Potential off-target genes are in yellow.

In addition to qPCR, RNAseq was performed using the other half of the total RNA extracted as described above, to determine the effects of *eve* and *hh* overexpression on other related developmental genes as well as screen for possible off-targeting (**Fig. 2B, Tables S1-S6**). Genome-wide transcriptome analysis confirmed the qPCR overexpression data. Higher TPM (Transcripts per million) values for both *eve* and *hh* were observed only in transheterozygous progeny for both crosses (**Table S2**). *Eve* has an abundance value of 91.52 TPM, a 5.7-fold increase (*P*= 3.56e-118) in transheterozygous PUb:dCas9-VPR/U6-sgRNA^*eve*^ eggs, significantly higher compared to 5.97 TPM and 1.45 TPM respectively for the control lines, PUb:dCas9-VPR and U6-sgRNA^*eve*^ (F_4,9_=66.45; *P*= 1.11e-06; one-way ANOVA and Tukey’s multiple-comparison test) (**Tables S1-3, S5**). Furthermore, *hh* has an abundance value of 11.03 TPM in transheterozygous eggs, representing a 2.8 fold increase (*P* = 1.09e-11), significantly higher than the two parental lines with 5.73 TPM and 0.79 TPM (F_4,10_ = 23.65; *P* = 4.41e-05; one-way ANOVA and Tukey’s multiple-comparison test) (**Fig. 2B, Tables S1-2, S4, S6**). Additionally, *hh* transheterozygous eggs also have a higher *eve* abundance value of 38.42 TPM, 3.5-fold increase (*P* = 3.67e-16) in transheterozygous individuals compared to 5.97, 1.45 and 1.53 TPM of the control lines (*P* < 0.005) (**Fig. 2B**). Finally, we uploaded all the RNAseq data into the integrative genomics viewer (IGV) and observed considerable upregulation in the transheterozygous samples as compared to the controls for both *eve* (**Fig. S1**) and *hh* (**Fig. S2**). Taken together, these data indicate robust and programmable target gene transactivation in transheterozygotes using both qPCR and RNAseq methodologies.

### Overexpression of *eve* and *hh* generated additional transcriptomic changes

We performed a differential expression analysis using the RNAseq data to search for genome-wide effects that could result from target gene overexpression. Differential expression comparisons between transheterozygous PUb:dCas9-VPR/U6-sgRNA^*eve*^and control lines (PUb:dCas9-VPR or U6-sgRNA^eve^) identified 6488 and 7816 genes (∼43%) with expression level changes (FDR<0.05) (**Fig. 2C, Tables S3, S5, S7**). A total of 3737 and 3529 of those genes for each comparison demonstrated relative increases in transcript abundance of more than 2-fold (2384 and 2100 genes respectively per each comparison passed FDR < 0.05 threshold). Among the differentially expressed genes that showed an increase in abundance (LogFC > 2, FDR < 0.05), the most significantly upregulated genes included genes pertaining to the expression of fibrinogen (AAEL013506), centrin (AAEL006492), lipase (AAEL000686), and an unspecified protein orthologous to PIWI in *Aedes albopictus* (AAEL021425) (**Fig. 2B)**.

Differential transcript abundance profile comparisons between transheterozygous PUb:dCas9-VPR/U6-sgRNA^*hh*^ and control lines (PUb:dCas9-VPR and U6-sgRNA^hh^) identified 6570 and 7257 genes (about 41%) with transcript level changes (FDR < 0.05) (**Fig. 2B, Tables S4, S6-7**) that likely are associated with *hh* overexpression. Of those genes, 3573 and 3645 respectively for each comparison showed more than two-fold changes in abundance (2207 and 2220 genes, respectively per each comparison passed the FDR < 0.05 threshold). Among the most differentially expressed genes that showed an increase in abundance (LogFC > 2, FDR < 0.05), the most statistically-significant genes that were upregulated were chaperonin and steroid dehydrogenase. The most significant downregulated genes were ribosomal proteins from both the 40S and 60S subunits (**Fig. 2B**). Taken together these results support the conclusion that the effect of *eve* and *hh* overexpression results in genome-wide differential gene expression.

To determine if dCas9-VPR has off-target activation effects as described previously [62,63], we bioinformatically predicted potential off-target binding sites using an optimal CRISPR target finding software [64,65]. We identified 62 possible off-target sites for *eve* sgRNAs and 23 possible off-targets for *hh* sgRNAs (**Tables S8-9**). We then searched to see if any nearby gene was upregulated in our RNAseq data. Among 85 potential off-target sites for the *eve* or *hh* sgRNAs, only 8 sites correlated to *eve* sgRNAs, localized near genes that were differentially expressed in our analysis with LogFC > 2 and FDR < 0.05 (**Fig. 2B, Tables S8-9**). No potential off-target sites related to *hh* sgRNAs showed significant transactivation. Taken together these results confirm that ectopic overexpression of the genes *eve* and *hh* can generate further transcriptional misregulation of many other genes. Furthermore the probability of off-target transactivation with dCas9-VPR is likely low. However, it is important to acknowledge that confirmation of off-target effects through RNAseq data is difficult due to the inability of distinguishing between either the upregulation of non-target genes resulting from simply upregulating an important developmental gene (i.e. *eve* and *hh* overexpression) or direct off-targeting by dCas9 on its own.

### Transactivation results in lethality

We screened transheterozygous progeny for potential observable phenotypes induced by CRISPRa-mediated transactivation. We observed that transactivation of *eve* and *hh* resulted in varied rates of embryonic lethality (**Fig. 3A**,**B**). Furthermore, rates of embryonic lethality were also affected by the paternal or maternal lineage of either CRISPRa element. To calculate these rates we measured the proportion of transheterozygous progeny between heterozygous CRISPRa parents compared to expected derived from Mendelian inheritance (25% transheterozygous offspring). For example, when transactivating *eve*, crosses between heterozygous PUb:dCas9-VPR/+♂ X U6-sgRNA^*eve*^/+♀ resulted in an average of 12% (52% reduction from expected) of transheterozygous progeny, whereas the reciprocal cross (PUb:dCas9-VPR/+♀ X U6-sgRNA^*eve*^/+♂) resulted in an average of 21% (16% reduction from expected). Heterozygous PUb:dCas9-VPR♂/+ and U6-sgRNA^*hh*^ ♀/+ crosses resulted in an average of only 1% (96% reduction from expected) survival of trans heterozygous progeny. The reciprocal cross ((PUb:dCas9-VPR/+♀ X U6-sgRNA^*hh*^/+♂) resulted in an average of 11% (56% reduction from expected) transheterozygous progeny (**Fig. 3B**). As seen from these results, targeting both genes resulted in higher rates of lethality when dCas9-VPR was inherited paternally while sgRNAs were inherited maternally. Furthermore, upregulating *hh* expression resulted in a higher frequency of lethality then targeting *eve*. Given the role these two genes play in development it was no surprise that their upregulation resulted in lethality.

**Figure 3.**
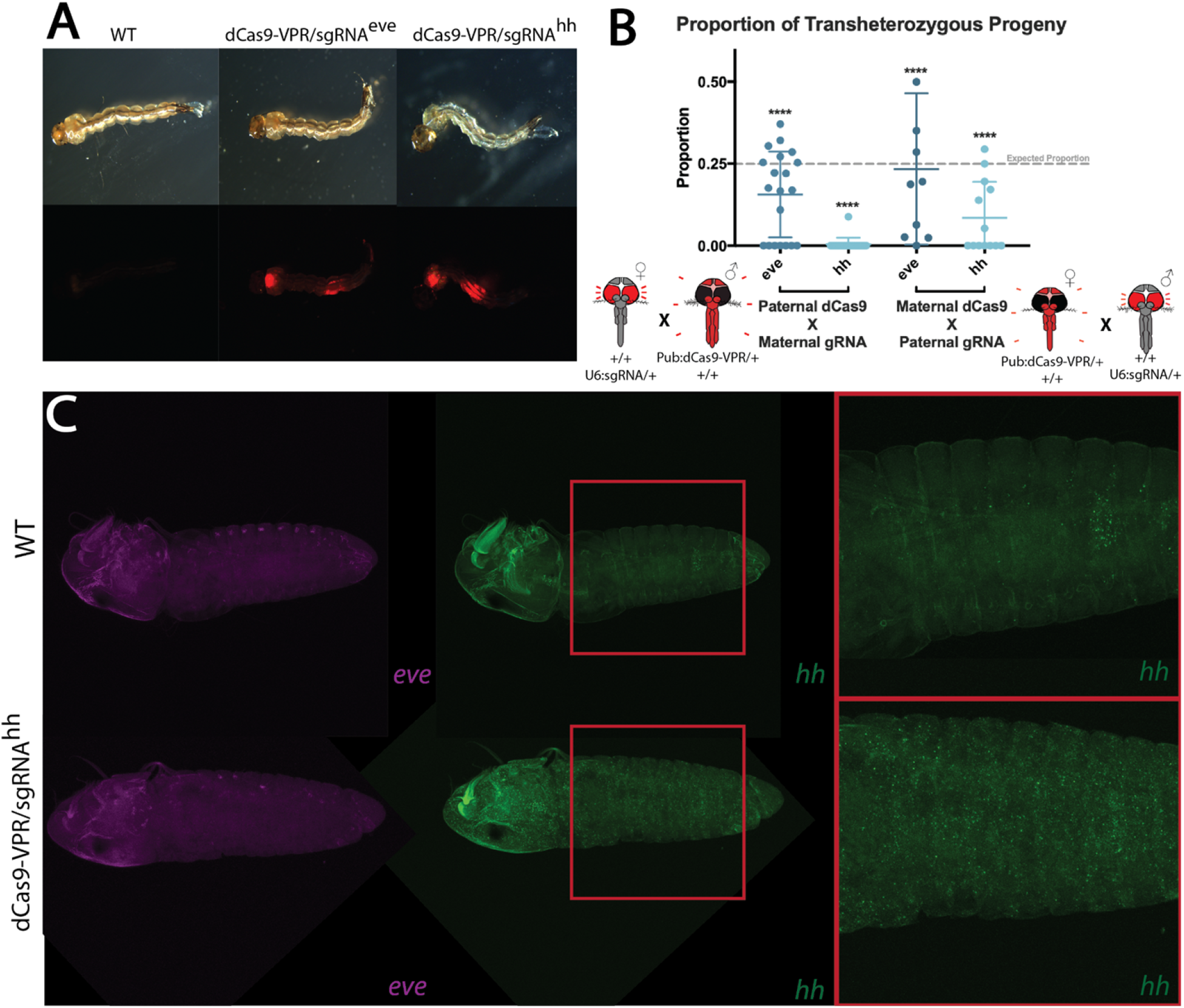
Phenotypic observations of CRISPRa mediated transactivation of *eve* and *hh*. **A)** Surviving transheterozygous progeny were collected and showed no significant external morphological differences to wild-type larvae. **B)** The proportions of transheterozygous progeny were calculated from single pair crosses between heterozygous dCas9-VPR and sgRNA parents. Rates of inheritance were compared to an expected rate of Mendelian inheritance, 25% transheterozygous individuals among total offspring, using a Chi-square test (\*\*\*\**p <* 0.0001). **C)** Further imaging of HCR *in situ* hybridization of *hh* probes showed upregulated and misregulated expression of *hh* compared to WT, while *eve* expression stayed relatively the same as that of WT.

We collected surviving transheterozygous larvae to screen them for visible morphological phenotypes that could be linked to ectopic expression of the target genes. Both PUb:dCas9-VPR/U6-sgRNA^*eve*^ and PUb:dCas9-VPR/U6-sgRNA^*hh*^ transheterozygotes did not display any observable differences in morphology compared to WT larvae, when observed under a stereoscope (**Fig. 3A**). We next wanted to look for observable overexpression in embryos. Therefore, we conducted *in situ* hybridization on PUb:dCas9-VPR/U6-sgRNA^*hh*^ embryos with probes designed for *eve* and *hh* transcripts. Imaging showed increased accumulation of *hh* transcripts throughout the embryo as well as a delayed developmental progress when compared to a WT embryo at the same developmental time point (**Fig. 3C)**. A significantly increased abundance of *eve* was not seen in PUb:dCas9-VPR/U6-sgRNA^*hh*^ embryos, however a subtle difference in localization was seen. This could be an artifact from *hh* transactivation affecting downstream genes. The ectopic expression of *hh* suggests that the PuB promoter can be used to transactivate within cells throughout the embryo. Furthermore, the delayed development of the PUb:dCas9-VPR/U6-sgRNA^*hh*^ embryo likely correlates with defects that would lead to the diminished numbers of transheterozygous progeny seen in our phenotypic screening crosses. Taken together, these results support the conclusion that our PUb:dCas9-VPR line is able to increase expression of target genes throughout the embryo.

### Overexpression of *AaRel1* results in virus suppression

The mosquitoToll and JAK-STAT immune signaling pathways and the siRNA immune pathway, play important roles in defending against arboviral infections [56,66,67] (reviewed in [68]). Here as proof of principle, we used CRISPRa to transactivate *AaRel1*, an NF-κB Relish-like transcription factor that mediates the Toll immune pathway’s antipathogenic action including the suppression of Dengue and Zika viruses [54–57]. First we generated a transgenic line expressing sgRNAs targeting the promoter region of *AaRel1* gene under the U6 promoter as described above (**Fig. 4A**) (U6:sgRNA^*rel1*^). To increase efficacy, the sgRNA line encodes four sgRNAs targeting four different regions within the promoter region of *AaRel1* (**Fig. 4A**). Specifically, the sgRNA target sites were designed from sequences within 0-250 base-pairs (bp) to the 5’-end of the transcription start site (TSS) to enable robust target gene transactivation. Expression of sgRNAs was ubiquitously-driven by either U6a or U6b promoters [61]. Fluorescent markers were encoded to confirm integration of the transgenes into the mosquito genome as well as tracking inheritance of the constructs. Two lines harboring the same sgRNA sequence but with different insertion sites were created and used for subsequent analysis U6:sgRNA^*rel1-A*^ and U6:sgRNA^*rel1-B*^

**Figure 4.**
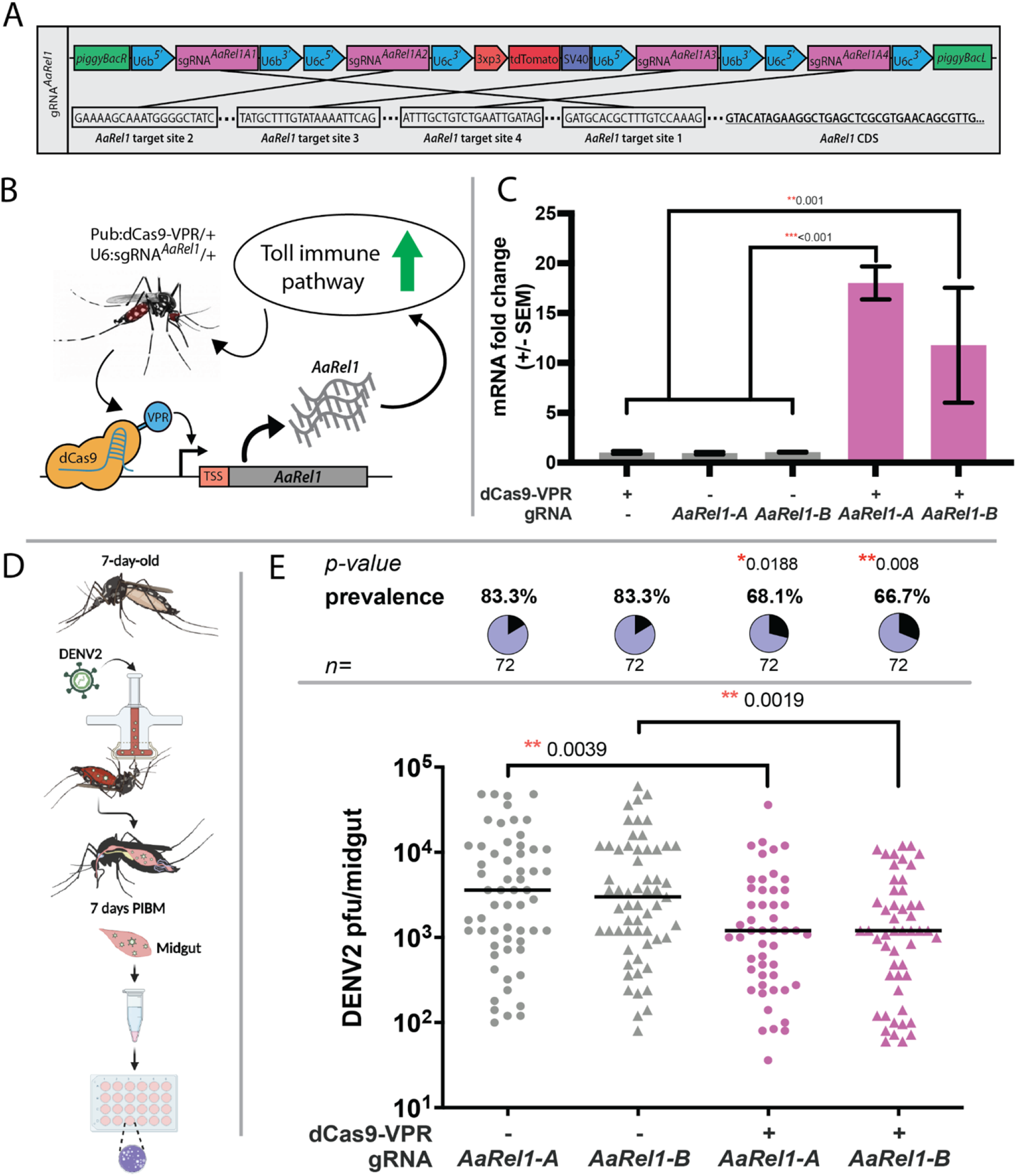
Generation of sgRNA-Rel1 lines for CRISPRa mediated transactivation of Toll immune pathway, RT-PCR validation of *AaRel1* overexpression and suppression of viral infection. **A)** Schematic representation of the sgRNA *AaRel1* construct used to create the U6:sgRNA-*AaRel1* lines. **B)** Schematic representation of the transgenic line expressing Pub:dCas9-VPR and the U6:sgRNA targeting the promoter region of *AaRel1* gene and inducing the activation of the Toll immune pathway. **C)** Quantification by qPCR of *AaRel1* gene expression in controls and transactivated lines. A one-way ANOVA and Tukey’s multiple-comparison test was performed between transheterozygous crosses and controls. \**P <* 0.05, \*\**P <* 0.01, \*\*\**P <* 0.001 **D)** Antiviral effect of CRISPRa mediated transactivation of *AaRel1* in the transheterozygous progeny. sgRNA-*AaRel1* line A and B (sgRNA-*AaRel1*-A and sgRNA-*AaRel1*-B, or named as U6:sgRNA^*rel1-A*^ and U6:sgRNA^*rel1-B*^) and transheterozygotes (dCas9-VPR/sgRNA-*AaRel1*-A and dCas9-VPR/sgRNA-*AaRel1*-B or named as dCas9-VPR/U6-sgRNA^*rel1-A*^ and dCas9-VPR/U6-sgRNA^*rel1-B*^) were orally infected with DENV2 as illustrated. **E)** The viral infection titers and infection prevalence of DENV2 were measured after midgut infection at 7 days post-infectious blood meal (PIBM). Plaque assays were used to determine viral titers and infection prevalence in individual mosquitoes with each dot representing the viral load from one midgut, and pie-chart indicating the infection prevalence. Horizontal black lines indicate the median of the viral loads. At least three replicates were pooled for the statistical analyses, using Mann-Whitney test to compare median virus titers and Fisher’s exact test to compare infection prevalence. \**P <* 0.05, \*\**P <* 0.01, *n*: total numbers of mosquitoes used in the assays. sgRNA-*AaRel1* lines were used as controls.

Subsequently, we performed crosses between PUb:dCas9-VPR/+ males and U6:sgRNA^*rel1-A*^/+ females and between between PUb:dCas9-VPR/+ males and U6:sgRNA^*rel1-B*^/+ females to quantify CRISPRa-mediated transactivation and its effect on virus replication. Resulting progeny were collected to quantify expression and transcript abundance of the target genes and perform Dengue virus challenge (**Fig. 4B**). We measured *AaRel1* transcript levels using quantitative real-time PCR (qPCR). Total RNA extracted from 1-day post-emergence adult transheterozygous females for dCas9-VPR/ U6-sgRNA^*rel1-A*^ and dCas9-VPR/U6-sgRNA^*rel1-B*^ and controls, including the parental lines (PUb:dCas9-VPR; U6-sgRNA^*rel1-A*^ U6-sgRNA^*rel1-B*^), were all used to measure transcript levels of *AaRel1*. qPCR analysis of *rel1* resulted in a significant 18.02 (± 0.95)-fold increase in transcript level for the transheterozygous females dCas9-VPR/U6-sgRNA^*rel1-A*^ and a significant 11.78 (± 3.33)-fold increase for the second transheterozygous line dCas9-VPR/U6-sgRNA^*rel1-B*^ (F**ig. 4C**) compared to the three controls (F_4,7_ = 637.6; *P* = 0.001; one-way ANOVA and Tukey’s multiple-comparison test) (**Fig. 4C**).

To investigate the impact of transactivation of *AaRel1* on dengue virus serotype 2 (DENV2) infection, we orally infected transheterozygous lines, dCas9-VPR/U6-sgRNA^*rel1-A*^ and dCas9-VPR/U6-sgRNA^*rel1-B*^, along with the siblings of the corresponding U6:sgRNA^*rel1-A*^ and U6:sgRNA^*rel1-B*^ mosquitoes as controls with an artificial blood meal containing 10^7^ PFU/mL virus particles using a glass membrane feeder system (**Fig. 4D**). The viral infection intensity and prevalence in the midgut at 7 days post-infectious blood meal (PIBM) was determined through plaque assay. The viral loads in the mosquito midguts were significantly reduced in the midgut tissues of both dCas9-VPR/U6-sgRNA^*rel1-A*^ and dCas9-VPR/U6-sgRNA^*rel1-B*^ lines when compared to the corresponding control mosquitoes, U6:sgRNA^*rel1-A*^ and U6:sgRNA^*rel1-B*^, with a significant 3-fold and 2.5-fold reduction in median virus titers, respectively (**Fig. 4E**, Mann-Whitney test, ** *P*<0.01). The infection prevalence also was reduced significantly with 18% and 20% reductions in both dCas9-VPR/U6-sgRNA^*rel1-A*^ and dCas9-VPR/U6-sgRNA^*rel1-B*^ lines, respectively, measured through Fisher’s exact test (**Fig. 4E**, \**P*<0.05, \*\**P*<0.01). The suppression of DENV2 viral infection in terms of both median intensity and prevalence indicates that CRISPRa-mediated transactivation of *AaRel1* in the midgut augmented the activity of the Toll pathway’s antiviral action in this tissue, consistent with previously published conclusion [56]. The extent of viral suppression is stronger than that displayed when using RNAi-mediated gene silencing of the Toll pathway negative regulator *Cactus*, most likely due to the more robust immune activation of the Toll immune signaling pathway achieved through CRISPRa-mediated transactivation of the Rel1 transcription factor.

## Discussion

Here we have demonstrated programmable transactivation of endogenous genes in *Ae. aegypti* using a CRISPRa system. As a proof of concept, we targeted conserved embryonic development genes, *eve* and *hh*. Utilizing both qPCR as well as RNAseq analysis we verified significant upregulation of the target genes resulting in increased transcript abundance only within individuals containing all CRISPRa components. In addition, we observed phenotypes related to overexpression of *eve* and *hh* in the form of embryonic lethality and transcript mispatterning in the embryo. It is not surprising that overexpression of these developmental genes resulted in embryonic lethality as similar phenotypes were observed in *D. melanogaster* using a CRISPRa approach [69].

The binary approach that maintains dCas9-VPR and sgRNAs in separate *Ae. aegypti* transgenic lines allow for further expansion and utility of the CRISPRa system. By separating these key components, a library of dCas9-VPR lines expressed under various promoters as well as sgRNAs targeting different genes can be designed and generated to allow for future flexibility. In this study, we designed a dCas9-VPR line under the polyubiquitin promoter (PUb), which is active in multiple cell types throughout all development as well as two lines expressing multiplexed sgRNAs targeting the promoter regions of developmental genes *eve* and *hh*. Additional sgRNA lines targeting other genes can be generated and crossed with our PUb:dCas9-VPR line or even additional dCas9-VPR lines under promoters that are spatially or temporally specific can be generated for more targeted transactivation of target genes.

Although we confirmed transactivation of target genes using our system, it is important to note that there is the inherent possibility of off-target effects. From our analysis we determined multiple possible off-target genes that resulted in an increase in expression within the RNAseq data. With the methods outlined in this paper, it is difficult to determine if the increased expression of these genes is linked to general effects of overexpressing our conserved target genes (*eve* or *hh*), or simply a result of direct off-targeting by the CRISPRa system. However to further understand and confirm transcriptome-wide effects caused from transactivating target genes, a variety of sgRNAs of different sequences could be used. Corroborating RNAseq results between these sgRNAs would strengthen the understanding of which affected non-target genes are differentially expressed due to target gene transactivation versus direct off-targeting. Furthermore, we transactivated two transcription factors known to play pivotal roles within embryonic developmental pathways. Transactivating genes that have no effect on transcription regulation may result in a reduced effect in differential expression within the transcriptome.

Overall, our results demonstrate that CRISPRa is viable in *Ae. aegypti* as a means to effectively transactivate specific genes. Current genetic engineering tools for *Ae. aegypti* have focused on knocking down, or out, genes as well as expressing transgenes. There has yet to be a tool for the targeted activation of endogenous genes. The ability to transactivate select endogenous genes can lend towards further understanding of *Ae. aegypti* genes through functional studies involving overexpression experiments as well further expanding the possibilities for vector control tools [70]. For example, a new class of vector control tools utilizing transactivation of endogenous genes can be developed. Previously, a promising system in *D. melanogaster* was used to generate synthetic speciation which could be used to introduce reproductive barriers within wild populations [71].

The dissection of potent roles of mosquito innate immunity requires powerful genetic tools. To investigate the role of immune signaling pathways, Toll, IMD, JAK-STAT, and siRNA immune pathway, previous studies rely on the tissue-specific overexpression of the key regulators in these pathways or RNAi-mediated silencing of either positive or negative regulators. The lack of specificity of RNAi-mediated gene silencing and flexibility and efficiency of tissue-specific transgenic overexpression of these key molecules have hampered the depth of the study of the immune signaling cascades and molecular mechanisms of the antiviral defenses in the mosquito’s innate immunity. Here as proof of principle, we have demonstrated that CRISPRa-mediated transcription activation of immune factors can be applied to study the role of immune pathway genes in the antiviral defenses. The advantage of this genetic tool is the activation of the key immune transcriptional factors results in restricting the viral infection in the mosquitoes more efficiently. With the rapid development of the tissue-specific dCas9-VPR lines in combinations with targets-specific sgRNA lines in the future, these key immune factors can be studied specifically in a spatio-temporal manner. This tool can be used to boost mosquitoes’ immunity against viral infections by transactivating genes involved in various immune pathways, such as Relish-like transcription factors Rel1 and Rel2, the two key downstream regulators of the Toll and IMD immune pathways [54,72,73], or the insect cytokine-like factor Vago [74,75], or DOME and HOP in the JAK-STAT pathway [67], or Dicer2 and R2d2 in the siRNA immune pathway [66]. The knowledge gained from these basic studies will further strengthen the development of vector control strategies. Further, this system could also be applied to *Ae. aegypti* to drive select genes into a population.

## Methods

### Mosquito rearing and maintenance

All *Ae. aegypti* lines used in this study were generated from the Liverpool strain. Colonies were reared at 27.0°C, 20–40% humidity, and a 12-h light/dark cycle. Adults were fed 0.3M aqueous sucrose *ad libitum*. To produce eggs, mature females were blood-fed using anesthetized mice. Oviposition cups were provided ∼3 days post blood-meal and eggs were collected and aged for ∼4 days before hatching. Matured eggs were submerged under deionized H_2_O and placed into a vacuum chamber set to 20 in Hg overnight. Emerged larvae were reared in plastic containers (Sterilite) with ∼3 liters of deionized H_2_O and fed daily with fish food (Tetramin). *Aedes* mosquitoes rearing at the Johns Hopkins Insectary Core Facility followed established standard procedures and were maintained on 10% sucrose solution under standard insectary conditions at 27±0.5°C and 75-80% humidity with a day:night light cycle of 14:10 h.

### Construct design and assembly

The Gibson enzymatic assembly method was used to engineer all constructs in this study. To generate the dCas9-VPR expressing construct, OA-986F (Addgene plasmid #183993), Our previously published OA-986A plasmid was used as a backbone [76]. Restriction enzymes, NotI and PmeI, were used to cut the plasmid backbone. DNA fragments containing the PUb promoter was amplified from the Addgene plasmid #100581 using primers 874r32 and 986F.C1. In addition, we generated 2 sgRNA plasmids each harboring 2 distinct sgRNAs targeting the promoter regions of *eve* (AAEL007369, OA-1053A, Addgene plasmid #184006) and *hh* (AAEL006708, OA-1053B, Addgene plasmid #184007). To engineer these plasmids, we modified plasmid OA-984 [77] (Addgene plasmid #120363) to contain 2 sgRNA sequences targeting either *eve* or *hh* driven by U6 promoters (sgRNA^*eve*^ and sgRNA^*hh*^). Restriction enzymes AvrII and AscI were used to create the plasmid backbone. Two GenPart fragments were synthesized from GenScript for each plasmid (OA-1053A or OA-1053B), one containing sgRNA1 driven by U6b (AAEL017774), while the other containing sgRNA2 driven by U6c (AAEL017763). For virus targeting, we generated one plasmid harboring 4 distinct sgRNAs targeting the promoter region of *AaRel1* (AAEL007696, OA-1127B, Addgene plasmid #190997). Firstly, two intermediate plasmids, OA-1127B.X1 (*rel1*-gRNA1&2) and 1127B.X2 (*rel1*-gRNA3&4), each harboring two gRNAs, were generated by cutting the same previous backbone plasmid OA-984 (Addgene plasmid #120363), with the restriction enzymes AvrII and AscI, and cloning in two GenPart fragments, which were synthesized from GenScript containing two gRNAs driven by U6b (AAEL017774) and U6c (AAEL017763) promoters respectively. Then, plasmid OA-1127B.X1 was linearized with the restriction enzyme FseI, and the insertion of U6b-*rel1*-gRNA3-U6c-*rel1*-gRNA4 was amplified with primers 1167.C5 and 1067.C6 from the plasmid OA-1127B.X2. During each cloning step, single colonies were chosen and cultured in Luria Bertani (LB) Broth with ampicillin. Plasmids were extracted using the Zippy plasmid miniprep kit ((Zymo Research, Cat No./ID: D4036) and sanger sequenced. Final plasmids were maxipreped using the ZymoPURE II Plasmid Maxiprep kit (Zymo Research, Cat No./ID: D4202) and sanger sequenced in preparation for embryonic microinjection. All primers are listed in **Table S10**. Complete plasmid sequences and plasmid DNA are available at www.addgene.com with accession numbers (#183993, #100581, #184006, #184007, #120363, #190997).

### Generation of transgenic lines

Transgenic lines were generated by microinjecting 0.5–1 h old pre blastoderm stage embryos with a mixture of the piggyBac plasmid (200 ng/µl) and a transposase helper plasmid (phsp-Pbac, (200 ng/µl)[78]. Embryonic collection and microinjections were performed following previously established procedures. After 4 days post-microinjection, G_0_ embryos were hatched in deionized H_2_0 under vacuum (20 in Hg). Emerged larvae were reared to pupal stage using previously established procedures. Surviving G_0_ pupae were sex-separated into ♀ or ♂ cages. WT ♀ or ♂ of similar age were added to cages of the opposite sex at a 5:1 ratio (WT:G_0_). Several days post-eclosion (∼4–7), a blood-meal was provided, and eggs were collected, aged, then hatched. Hatched larvae were screened and sorted for expression of relevant fluorescent markers using a fluorescent stereo microscope (Leica M165FC). Each individual line was maintained as mixtures of homozygotes and heterozygotes with periodic selective elimination of wild-types.

### Generating and screening for CRISPRa transheterozygotes

To test for transactivation PUb-dCas9 females with males from our sgRNA lines, after allowing the crosses to mate for 3 days, females were blood-fed for 2 consecutive days. Three days after blood-feeding, females were individually captured in plastic vials lined with moistened paper. Captured females were kept for 2 days to allow for egg laying and removed afterwards. Collected eggs were either processed for RNA collection, fixed and dechorionated for staining, or hatched to screen surviving progeny.

### Total RNA collection and sequencing

To directly observe and quantify targeted *eve* and *hh* transactivation mediated by PUb:dCas9-VPR, we collected transheterozygous embryos for RNA extraction and subsequent sequencing. Embryos were collected 24 h post-oviposition from F1 transheterozygous lines (PUb:dCas9-VPR/U6-sgRNA^*eve*^ and PUb:dCas9-VPR/U6-sgRNA^*hh*^) as well as parental lines (PUb:dCas9-VPR; U6-sgRNA^*eve*^ and U6-sgRNA^*hh*^). Three biological replicates per line were collected for a total of 15 samples. Additionally, To quantify the transactivation of *rel1*, one day old adult females were used for RNA extraction and subsequent qPCR analysis. Three biological replicates per line were collected for a total of 15 samples, including the two transheterozygous lines (PUb:dCas9-VPR/U6-sgRNA^*rel1-A*,^and PUb:dCas9-VPR/U6-sgRNA^*rel1-B*^) and the three parental controls (PUb:dCas9-VPR, U6-sgRNA^*rel1-A*^ and U6-sgRNA^*rel1-B*^). Total RNA was extracted using a Qiagen RNeasy Mini Kit (Qiagen 74104). Following extraction, total RNA was treated with an Invitrogen DNase treatment kit (Invitrogen AM1906). RNA concentration was analyzed using a Nanodrop OneC UV-vis spectrophotometer (ThermoFisher NDONEC-W). About 1 μg of RNA was used to synthesize cDNA with a RevertAid H Minus First Strand cDNA Synthesis kit (Thermo Scientific). CDNA was diluted 50 times before use in Real-Time quantitative PCR (RT-qPCR). RT-qPCR was performed with SYBR green (qPCRBIO SyGreen Blue Mix Separate-ROX Cat #: 17-507B, Genesee Scientific). 4 μl of diluted cDNA was used for each 20 μl reaction containing a final primer concentration of 200 nM and 10 μl of SYBR green buffer solution. Three technical replicates for each reaction were performed to correct for pipetting errors. The following qPCR profile was used on the LightCycler® instrument (Roche): 3 min of activation phase at 95°C, 40 cycles of 5 s at 95°C, 30 s at 60°C. **Table S10** lists the primers for *eve, hh, AaRel1* and *rpl32* (*ribosomal protein L32*). The *rpl32* gene was used as a reference gene [79] to calculate relative expression level of *eve, hh* and *AaRel1* with the manufacturer software and the delta-delta Ct method (2^−ΔΔCt^). Difference in expression of *eve, hh* and *AaRel1* between controls and transactivated lines, was statistically tested with one way ANOVA and a Tukey’s multiple-comparison test in RStudio statistical software (version 1.2.5033, © 2009-2019).

Collected RNA for *eve* and *hh* was also used to perform RNA-seq analyses in order to further validate results from qPCR analysis as well as detect other genes affected by *eve* and *hh* upregulation and potential off target genes. RNA integrity was assessed using RNA 6000 Pico Kit for Bioanalyzer (Agilent Technologies 5067-1513) and RNA-seq libraries were constructed using NEBNext Ultra II RNA Library Prep Kit for Illumina (NEB E7770) following manufacturer’s instructions. Libraries were sequenced on Illumina HiSeq2500 in single read mode with the read length of 50 nt and sequencing depth of 20 million reads per library. Basecalling was performed with RTA 1.18.64 followed by conversion to FASTQ with bcl2fastq 1.8.4.

### Bioinformatics: Quantification and differential expression analysis

RNA integrity was assessed using the RNA 6000 Pico Kit for Bioanalyzer (Agilent Technologies #5067-1513), and mRNA was isolated from ∼1 μg of total RNA using NEBNext Poly(A) mRNA Magnetic Isolation Module (NEB #E7490). RNA-seq libraries were constructed using the NEBNext Ultra II RNA Library Prep Kit for Illumina (NEB #E7770) following the manufacturer’s instructions. Libraries were quantified using a Qubit dsDNA HS Kit (ThermoFisher Scientific #Q32854), and the size distribution was confirmed using a High Sensitivity DNA Kit for Bioanalyzer (Agilent Technologies #5067-4626). Libraries were sequenced on Illumina HiSeq2500 in single read mode with the read length of 50 nt and sequencing depth of 20 million reads per library. Basecalling was performed with RTA 1.18.64 followed by conversion to FASTQ with bcl2fastq 1.8.4. The reads were mapped to *Aedes aegypti* genome AaegL5.0 (GCF_002204515.2) supplemented with PUb-dcas9 transgene sequence using STAR [80]. Gene expression was then quantified using featureCounts against NCBI *Aedes aegypti* Annotation Release 101 (GCF_002204515.2_AaegL5.0_genomic.gtf). TPM values were calculated from counts produced by featureCounts and combined (combined_count_tpm.aaegl5_dCas9-vpr.xlsx). Hierarchical clustering and PCA analyses were performed in R and plotted using R package ggplot2. Differential expression analyses between controls (PUb:dCas9-VPR; U6-sgRNA^*eve*^ and U6-sgRNA^*h*^) and transheterozygous lines (PUb:dCas9-VPR/U6-sgRNA^*eve*^ and PUb:dCas9-VPR/U6-sgRNA^*hh*^) were performed with DESeq2 (deseq2_sgRNA_transhet_Eve.xlsx, deseq2_dCAS9_VPR_transhet_Eve.xlsx, deseq2_sgRNA_transhet_hh.xlsx, deseq2_dCAS9_VPR_transhet_hh.xlsx). Illumina RNA sequencing data has been deposited to the NCBI-SRA, (PRJNA851480, reviewer link: https://dataview.ncbi.nlm.nih.gov/object/PRJNA851480?reviewer=pslogavc8iqjd3qnail9dfk0nb).

### Phenotypic Screening

To collect egg lays from single pair mating events, female and male mosquitoes were allowed to mate for 3 days post eclosion. Females were given a blood meal for 2 consecutive days. The day following the 2nd blood meal, blood-fed females were placed independently into plastic drosophila vials lined with wet paper and plugged with a foam plug. The females were kept in the vials for 2–3 days to allow for egg laying. Following oviposition onto the paper lining the drosophila vial, females were released into a small cage and egg lays were collected, counted, and allowed to mature to full development (∼4 days) in their original vials. Matured eggs were hatched within their original vial under vacuum overnight. Following hatching, egg papers were removed from the vials to allow for more space for the larvae to grow. At the L3 stage, progeny were screened, scored, and counted for expression of opie-2-dsRed and 3xP3-tdTomato using a fluorescent stereoscope (Leica M165FC). The difference in total larval counts compared to total egg counts were considered to be dead during embryonic or early larval stages. Surviving transheterozygous individuals were collected for further observation and analysis.

### *In situ* hybridization and embryo imaging

Embryos 24 h post-oviposition were collected, fixed, and dechorionated using previously described methods [81]. To more effectively remove the endochorion, peeling was performed in a mixture of PBS/PBT instead of methanol/ethanol. To free the embryo from the endochorion, fine tip forceps were used to crack a ring around the middle of the egg, taking care to not puncture the embryo. Cracked embryos were then briefly placed in methanol then back into PBS to improve detachment of the embryo from the endochorion. Each end of the endochorion was then teased off of the embryo. Yolk clarification was then performed according to previously described methods [82]. HCR *in situ* hybridization was performed using previously described methods [83]. HCR probes purchased from Molecular Instruments. Stained embryos were imaged using a Leica SP8 Confocal with Lightning Deconvolution.

### Virus propagation and oral viral infections in *Ae. aegypti*

DENV serotype 2 New Guinea C strain (DENV2) was cultured in *Aedes albopictus* C6/36 cells (ATCC CRL-1660), and viral stocks were prepared as previously described in [66,67,84]. All infection procedures were performed under BSL2 conditions for DENV2. Briefly, C6/36 cells were cultured in MEM medium (Gibco, Thermo Fisher Scientific, USA) supplemented with 10% heat-inactivated fetal bovine serum (FBS), 1% penicillin-streptomycin, and 1% non-essential amino acids and maintained in a tissue culture incubator at 32°C and 5% CO_2_. Baby hamster kidney strain 21 (BHK-21, ATCC CCL-10) cells were maintained at 37°C and 5% CO_2_ in the DMEM medium (Gibco, Thermo Fisher Scientific) supplemented with 10% fetal bovine serum (FBS), 1% penicillin-streptomycin, and 5µg/ml Plasmocin (InvivoGen, USA). For the preparation of DENV2 viral stocks, C6/36 cells grown to 80% confluence were infected with DENV2 at a multiplicity of infection (MOI) of 10 and incubated at 32°C and 5% CO_2_ for 5∼6 days. Virus was harvested by three freeze-thaw cycles using dry ice and a water bath (37°C), followed by centrifugation at 2,000 rpm for 10 min at 4°C. The supernatant from this cell lysis was mixed with the original cell culture supernatant to yield the final viral stock. Viral stocks were aliquoted and stored at -80°C for long-term storage.

Seven-day-old mosquitoes were orally infected with DENV2 through artificial glass membrane feeders as previously described [66,67,84]. A portion of each blood meal was frozen, and back titrated by plaque assay on the BHK-21 cells at 37°C. Mosquitoes were starved for 24 h prior to being offered the blood meal and were allowed to feed for ∼30 min. Fully engorged mosquitoes were sorted into soup cups, with no more than 60 individuals per cup. Each experiment was performed in at least three biological replicates, as indicated.

### Plaque assays for viral titration

DENV2 infected mosquito samples were titrated in the BHK-21 cell culture and plaque assays were used to determine infection prevalence and the viral titers. In brief, mosquito midguts were collected at 7 days post-infectious blood meal (PIBM) in 150 µl of complete DMEM medium with glass beads. A Bullet Blender (Next Advance, USA) was used to homogenize the tissue samples, and serial dilutions were prepared with DMEM complete medium. The BHK-21 cells were split to give a 1:10 dilution and grown on 24-well plates to 80% confluence 1-2 days before the plaque assays. After serially diluted, the mosquito tissue or viral stock samples (100 µl each) were added to the BHK-21 cells, followed by incubation at room temperature for 15 min on a rocking shaker (VWR International LLC) and subsequent incubation at 37°C with 5% CO_2_ in a cell incubator (Thermo Fisher Scientific) for another 45 min. The 24-well plates with infected BHK-21 cells were overlaid with 1 ml of 0.8% methylcellulose in complete DMEM medium with 2% FBS and incubated for 5 days in a cell culture incubator (Thermo Fisher Scientific, 37°C and 5% CO_2_). Plaques were fixed and developed with staining reagent (1% crystal violet in 1:1 methanol/acetone solution) at room temperature for 2 h. Plates were rinsed with distilled water and air-dried, and plaques were counted and multiplied by the corresponding dilution factors to calculate the plaque-forming units (PFUs) per sample. Three biological replicates were done with viral infection assays, and three replicates were pooled to generate the final figure. Dot-plot of infection intensities and pie-chart of infection prevalence were prepared with GraphPad Prism 9 software, and the significance of the infection intensities was determined by Mann-Whitney test and infection prevalence by Fisher’s exact test.

## Data Availability

All plasmids and annotated DNA sequence maps are available at www.addgene.com under accession numbers: #183993, #100581, #184006, #184007, #120363. Raw sequencing data are available at NCBI Sequence Read Archive, accession number PRJNA851480 (reviewer link: https://dataview.ncbi.nlm.nih.gov/object/PRJNA851480?reviewer=pslogavc8iqjd3qnail9dfk0nb).

## Ethical conduct of research

All animals were handled in accordance with the Guide for the Care and Use of Laboratory Animals as recommended by the National Institutes of Health and approved by the UCSD Institutional Animal Care and Use Committee (IACUC, Animal Use Protocol #S17187) and UCSD Biological Use Authorization (BUA #R2401). The protocol (permit # MO15H144) was approved by the Animal Care and Use Committee of Johns Hopkins University. Commercially obtained anonymous human blood type O+, and untyped human serum (InterState Blood Bank, Inc.) were used for DENV2 infection assays in the mosquitoes, and informed consent was therefore not required.

## Acknowledgements

We thank Judy Ishikawa for mosquito husbandry assistance. We thank the Johns Hopkins Malaria Research Institute Insectary for providing the mosquito-rearing facility and the Parasitology Core facilities for providing the naïve human blood. Figures were Created with BioRender.com. This work was supported by funding from a DARPA Safe Genes Program Grant (HR0011-17-2-0047), and NIH awards (R01AI151004, DP2AI152071, and R21AI149161) awarded to O.S.A, and NIH award R01AI141532 awarded to G.D. The views, opinions, and/or findings expressed are those of the authors and should not be interpreted as representing the official views or policies of the U.S. government. AAJ is a Donald Bren Professor at the University of California, Irvine.

## Author Contributions

O.S.A., A.A.J., and G.D. conceived and designed the experiments. M.B., E.D.B., Y.D., Y.Z., M.L., T.Y., A.B., and V.B. performed molecular and genetic experiments. I.A. performed the RNA sequencing experiments and analysis. M.W.P and Y.Z., performed *in situ* hybridization experiments. YD performed viral infection experiments. All authors contributed to the writing, analyzed the data, and approved the final manuscript.

## Competing Interests

O.S.A is a founder of both Agragene, Inc. and Synvect, Inc. with equity interest. The terms of this arrangement have been reviewed and approved by the University of California, San Diego in accordance with its conflict of interest policies. All other authors declare no competing interests.

---

**Figure S1.**
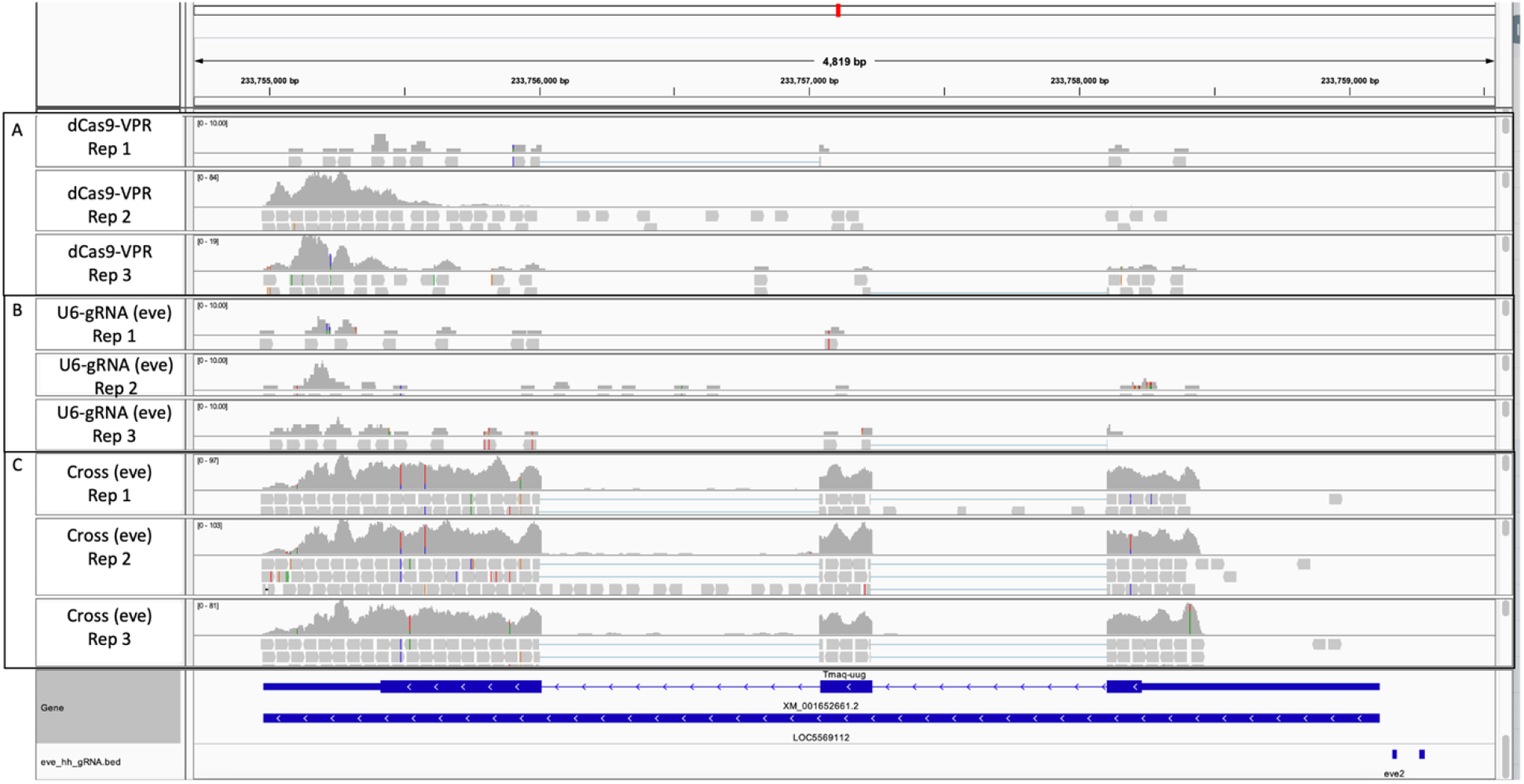
Integrative Genomics Viewer (IGV) Snapshot of the RNAseq data for *eve* overexpression. **A)** Three RNAseq replicates of dCas9-VPR. **B)** Three RNAseq replicates of U6-gRNA (*eve*). **C)** Three RNAseq replicates of the cross between dCas9-VPR and U6-gRNA. The gene structure of *eve* and two *eve* gRNA binding sites are indicated in blue at the bottom.

**Figure S2.**
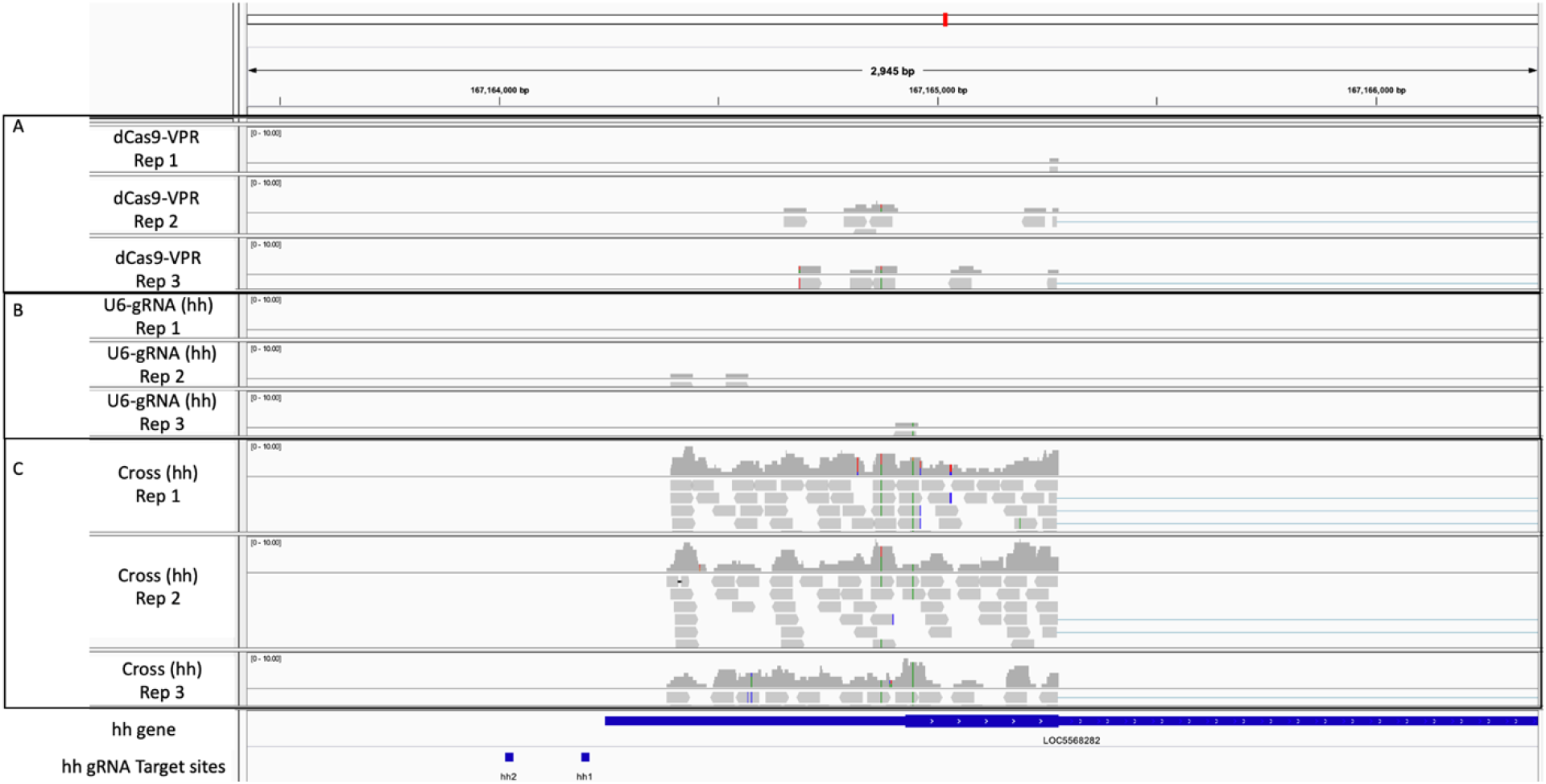
Integrative Genomics Viewer (IGV) Snapshot of the RNAseq data for *hh* overexpression. **A)** Three RNAseq replicates of dCas9-VPR. **B)** Three RNAseq replicates of U6-gRNA (*hh*). **C)** Three RNAseq replicates of the cross between dCas9-VPR and U6-gRNA. The gene structure of *hh* and two *hh* gRNA binding sites are indicated in blue at the bottom.

